# Unmatched cell line collections are not optimal for identification of PARP inhibitor response and drug synergies

**DOI:** 10.1101/2025.04.28.650908

**Authors:** Zoe Phan, Kristine J. Fernandez, C. Elizabeth Caldon

## Abstract

**Purpose:** While PARP inhibitors have shown great efficacy in patients with *BRCA1* and *BRCA2* mutations, many preclinical studies, including combination therapies, fail to translate into clinical approval, potentially due to the limitations of preclinical models. The goal of this brief report was to identify appropriate cell line models to investigate PARP inhibitor sensitivity and synergies.

**Methods:** An *in-silico* study of cell line collections was performed to assess the correlation between *BRCA1* or *BRCA2* mutations and sensitivity to PARP inhibitor monotherapy or combination therapy with a platinum-based chemotherapy. Subsequently, we characterised an isogenic model with matched cell lines containing *Brca1* and *Brca2* mutations and investigated the response of cells to PARP inhibitor alone, or in combination with chemotherapy.

**Results:** Using cell line collections, cell lines with *BRCA1* and *BRCA2* alterations are not associated with increased sensitivity to PARP inhibitors. Other factors, such as high *PARP1* expression and low-level genome alterations showed correlation with increased sensitivity to a PARP inhibitor. Furthermore, cell line collections did not reflect the synergy observed in patients to combination PARP inhibitor and platinum-based chemotherapy. Subsequently, we demonstrated that the ID8 isogenic model cell lines with specific mutations in *Brca1* and *Brca2* represent patient response to PARP inhibitor monotherapy, and in combination with chemotherapy.

**Conclusions:** This study suggests exercising caution when using cell line collections as part of model selection when investigating PARP inhibitor sensitivity and synergy. Our data proposes that using an isogenic preclinical model is more likely to accurately reflect patient response.

---

In the clinic, patients with a *BRCA1* or *BRCA2* mutation derive a significant benefit with PARP inhibitor treatment, a finding which has been confirmed in dozens of clinical trials across multiple cancer types including ovarian, breast and prostate cancers(1-3). Given the current success, but limited use, of PARP inhibitors, appropriate pre-clinical models are required to develop new applications for PARP inhibitors. Of particular difficulty is extending the use of PARP inhibitors to identify where they can successfully be used in combination with other therapies. While there have been many combination therapy trials and preclinical studies, there are only a handful of approvals (eg talazoparib and enzalutamide in metastatic prostate cancer(4), and bevacizumab and olaparib in ovarian cancer(5)). The failure of many preclinical studies to translate to clinical approval is potentially due to the limitations of current preclinical models(6). Therefore, in this brief report, we aimed to identify appropriate cell line models suitable for both PARP inhibitor sensitivity and synergy studies.

Firstly, we assessed the benefit of cell line collections in signal seeking PARP inhibitor studies by systematically analyzing cell lines from the Cancer Cell Line Encyclopedia (Broad, 2019)(7-9), accessed via cBioPortal. We extracted information on *BRCA1* and *BRCA2* mRNA expression, mutation status, and copy number alterations, alongside IC_50_ values for PARP inhibitor monotherapies. The list of cell lines alongside *BRCA1* and *BRCA2* mutation status are found in Supplementary Tables 1 and 2.

Surprisingly, in *BRCA1*-mutant cell lines, *BRCA1* mRNA expression showed no correlation with IC_50_ response to olaparib (r = -0.27, p = 0.42) (Figure 1A, left). Counterintuitively, among *BRCA1*-wildtype cell lines, lower *BRCA1* mRNA expression was significantly associated with reduced sensitivity to olaparib, as indicated by the significant negative correlation (r = -0.20, p < 0.0001) (Figure 1A; left). This was unexpected, given that *BRCA1*-mutant cell lines exhibited lower *BRCA1* mRNA expression than *BRCA1-*wildtype cell lines (Supplementary Figure 1A).

**Figure 1.**
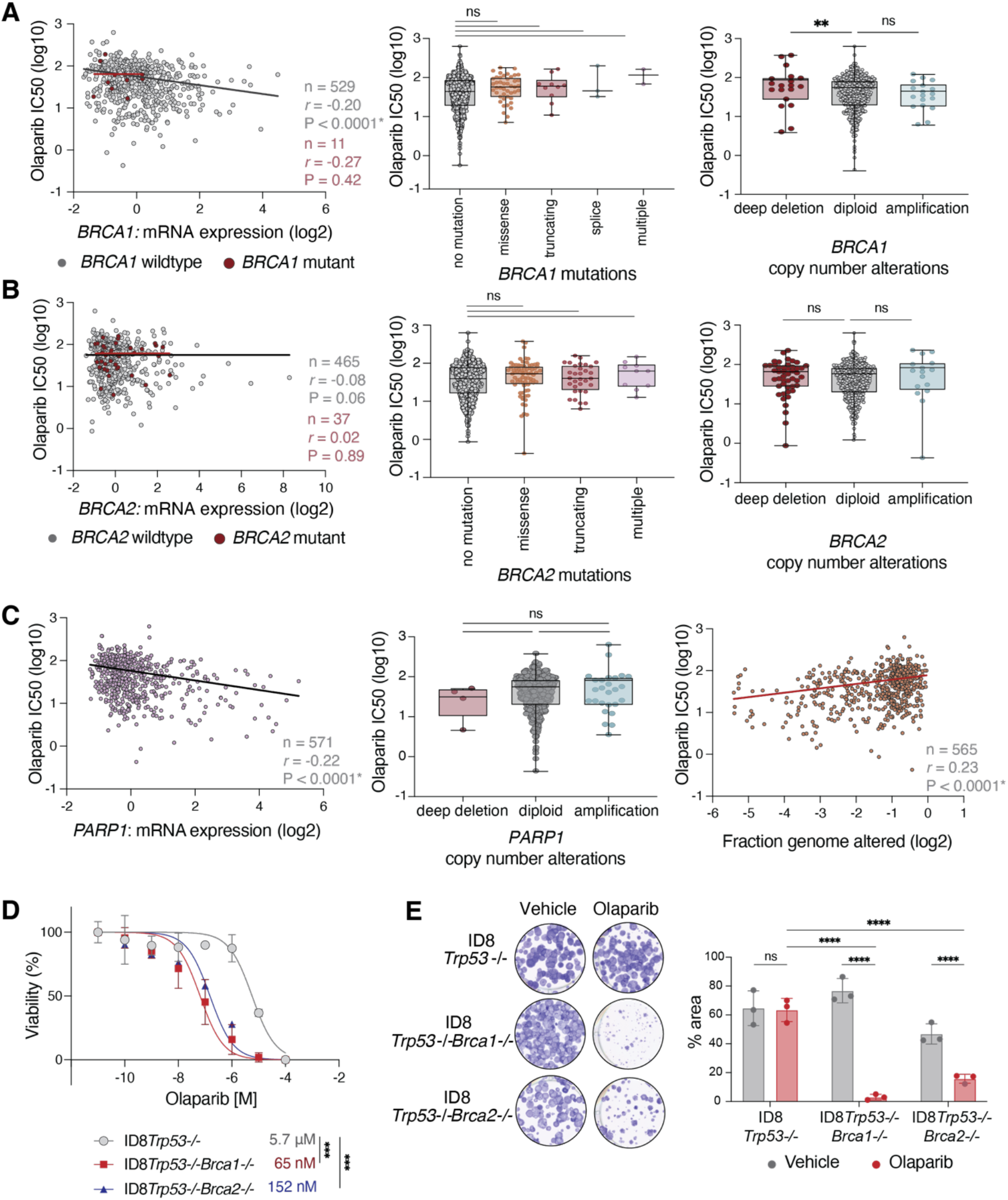
Model selection is critical for detecting changes in PARP inhibitor sensitivity due to *BRCA1/BRCA2* mutations. **(A-B)** Analysis of cell line databases demonstrates that **(A)** *BRCA1*-mutant and **(B)** *BRCA2*-mutant cell lines across different cancer types do not fully represent patient responses to olaparib. Left panels: Correlation between olaparib IC_50_ response and *BRCA1/BRCA2* mRNA expression relative to diploid samples. Red dots indicate cell lines with predicted driver mutations in *BRCA1* (n = 11) and *BRCA2* (n = 37), while grey dots indicate wildtype *BRCA1* (n = 529) and *BRCA2* (n = 465) cell lines. Best-fit linear regressions (red and black lines) illustrate positive or negative correlations. Middle panels: Olaparib IC_50_ response in *BRCA1-* and *BRCA2-*mutant cell lines with different mutation types: no mutation (n = 810 for *BRCA1*, n = 752 for *BRCA2*), missense (n = 45, n = 76), truncating (n = 10, n = 32), splice (n = 3) and multiple (n = 2, n = 10). Right panels: Olaparib IC_50_ response in cell lines with different *BRCA1/BRCA2* copy number alteration statuses. IC_50_ values are shown for cell lines classified as deep deletion (n = 19 for *BRCA1*, n = 47 for *BRCA2*), diploid (n = 518, n = 492), and amplification (n = 18, n = 16). **(C)** High *PARP1* expression and low genome alterations are associated with increased olaparib sensitivity in cell line collections. Left panel: Correlation between olaparib IC_50_ response and *PARP1* mRNA expression relative to diploid samples (n = 571). Middle panel: Olaparib IC_50_ response in cell lines with *PARP1* copy number alterations classified as deep deletion (n = 4), diploid (n = 525) and amplification (n = 26). Right panel: Correlation between olaparib IC_50_ response and fraction genome alterations (n = 565). Statistical analyses were determined by two-sided Pearson’s correlation test (left panels) and one-way ANOVA (middle and right panels). Data were accessed via cBioPortal. **(D-E)** ID8 isogenic models are optimal for investigating PARP inhibitor sensitivity. **(D)** Cell viability assays demonstrate that ID8*Trp53-/-Brca1-/-* and ID8*Trp53-/-Brca2-/-* cells are more sensitive to olaparib compared to ID8 *Brca* wildtype cells. Cells were treated with a range of olaparib doses for 3 days and IC_50_ values were determined using alamarBlue assay (n = 3 biological replicates). Some error bars are not visible as they are smaller than the data points. **(E)** Colony forming assays of ID8 cells treated with vehicle (DMSO) or 1 µM olaparib for 48 hours demonstrate a decrease in colony formation with olaparib treatment in *Brca*-mutated cell lines. Statistical analyses across cell lines were determined by two-way ANOVA. Statistical analyses for comparison within cell lines were determined by two-sided t-test. N = 3 biological replicates. ** p < 0.01; *** p < 0.001; **** p < 0.0001; ns = non-significant.

Next, we examined whether specific mutation types influence PARP inhibitor response. We found no significant difference in IC_50_ values between *BRCA1*-mutant cell lines (missense, truncating, splice or multiple mutations) and *BRCA1-*wildtype cell lines (Figure 1A, middle). Even cell lines harboring predicted driver mutations failed to exhibit increased sensitivity to PARP inhibitors compared to those without *BRCA1* mutations (Supplementary Figure 1C). Contrary to expectations, cell lines with a *BRCA1* deep deletion demonstrated reduced sensitivity to olaparib compared to those with normal *BRCA1* copy number (p = 0.01) (Figure 1A; right). This finding is particularly intriguing, given that *BRCA1* deep deletion is associated with lower *BRCA1* mRNA levels (Supplementary Figure 1B).

Analysis of *BRCA2*-alterated cell lines revealed similar trends to those observed in *BRCA1*-altered cell lines. In *BRCA2*-mutant cell lines, *BRCA2* mRNA expression did not correlate with olaparib response (r = 0.02, p = 0.89) (Figure 1B; left). In *BRCA2*-wildtype cell lines, there was a significant negative correlation between mRNA expression and treatment response (r = -0.08, p = 0.06) (Figure 1B; left). There was no association between *BRCA2* mutation type and olaparib response (Figure 1B; middle). Similarly, *BRCA2* deep deletion did not alter olaparib sensitivity compared to diploid cell lines, despite being associated with lower *BRCA2* mRNA expression (Figure 1B; right) (Supplementary Figure 1B).

We further analysed the impact of *BRCA1* and *BRCA2* alterations on the response to additional PARP inhibitors, including talazoparib, rucaparib, veliparib, as well as the platinum-based chemotherapy, cisplatin (Supplementary Figure 2 and 3). Across all analyses, *BRCA1-* or *BRCA2-*mutant or deep-deleted cell lines did not demonstrate increased sensitivity to PARP inhibitors or platinum-based chemotherapy. Furthermore, subgroup analyses using only ovarian cancer cell lines, representing the clinically most responsive group, showed no association between *BRCA1* and *BRCA2* alterations and response to olaparib (Supplementary Figure 4).

Given that *BRCA1* and *BRCA2* mutations in cell line collections did not predict PARP inhibitor response, we explored alternative biomarkers that may better correlate with sensitivity. Notably, high *PARP1* mRNA expression was significantly associated with increased responsiveness to olaparib across all cell lines (r = -0.22; p <0.0001) (Figure 1C; left). However, *PARP1* copy number alterations (deletion or amplification) did not impact olaparib response (Figure 1C; middle).

Another predictive factor was the fraction genome altered in each cell line, where cell lines with fewer genomic alterations were more sensitive to olaparib (r = 0.23, p < 0.0001) (Figure 1C; right). Together, these findings highlight that in cell lines, both high *PARP1* gene expression and low-level genome alterations are stronger predictors of olaparib response than *BRCA* alterations (Figure 1A/B versus C).

Given that cell line collections are not representative of *BRCA1-* and *BRCA2-*altered patient response to PARP inhibitors, we hypothesized that an isogenic model hosting specific gene alterations would be more appropriate to explore cellular responses to PARP inhibitor treatment. Therefore, we characterized the ID8 model of ovarian cancer, with matched cell lines with specific mutations in *Brca1* and *Brca2*(10, 11). ID8*Trp53-/-Brca1-/-* and ID8*Trp53-/-Brca2-/-* cells were significantly more sensitive to olaparib than ID8*Trp53-/-Brca-*wildtype cells by measuring cell viability (Figure 1D). Furthermore, cells with *Brca1* or *Brca2* mutation demonstrated a significant decrease in the ability to form colonies in the presence of olaparib compared to *Brca*-wildtype cells (Figure 1E). Thus, the ID8 model with specific mutations in *Brca1* and *Brca2* are significantly more sensitive to olaparib compared to their *Brca-* wildtype counterpart, in line with previous observations(10-12).

In a previous study, using clinical data, we demonstrated that *BRCA1/2*-mutated patients respond better to combination PARP inhibition and chemotherapy compared to *BRCA*-wildtype patients(6). Therefore, we assessed whether cell line collections with a *BRCA1/2* mutation recapitulates this response observed in the clinic by analyzing synergy data from the Genomics Drug Sensitivity in Cancer drug combination database (Wellcome Sanger Institute)(13). We examined the clinically effective combination of PARP inhibitor and platinum-based chemotherapy in cell lines with a predicted driver mutation in *BRCA1* or *BRCA2* compared to *BRCA1/BRCA2* wildtype cell lines(14-19). When investigating synergy between olaparib and cisplatin, there was no significant difference in ΔIC_50_ or ΔEMax based on mutation status (Figure 2A). Furthermore, these values did not cross ΔIC_50_ ≥ 3 and ΔEMax ≥ 0.2, and therefore, were deemed antagonistic.

**Figure 2.**
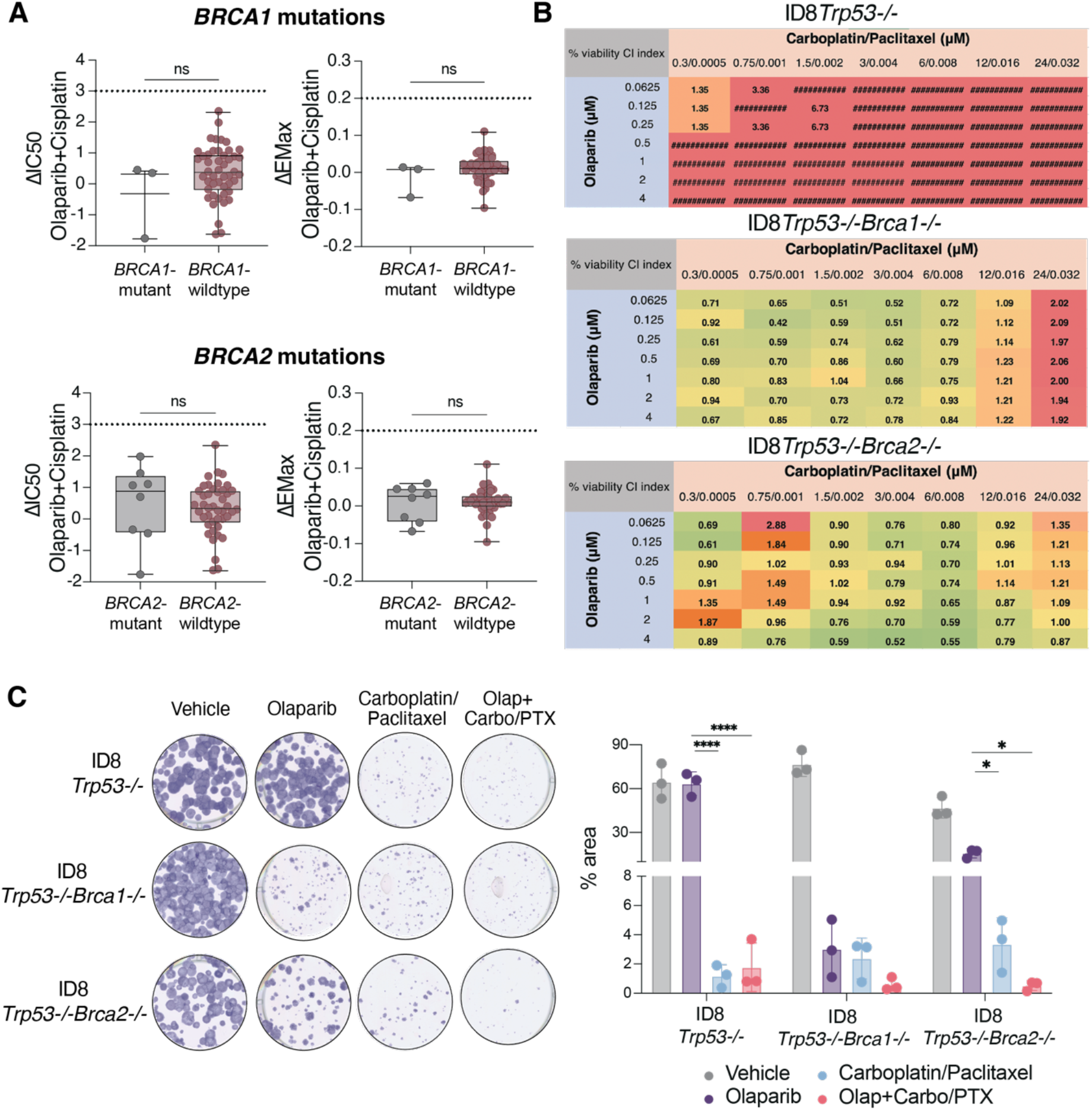
Isogenic models with specific *BRCA1/BRCA2* mutations are suitable for investigating PARP inhibitor drug synergies. **(A)** Analysis of breast cancer cell line collections shows no significant association between *BRCA1/BRCA2* mutation status and sensitivity to combined PARP inhibitor and chemotherapy compared to *BRCA*-wildtype. ΔIC_50_ and ΔEMax drug combination response for olaparib + cisplatin are shown for breast cancer cell lines with a *BRCA1* (n = 3) or *BRCA2* mutation (n = 8) and *BRCA-*wildtype (n = 47 for *BRCA1*, n = 43 for *BRCA2*). Synergism is defined as ΔEMax ≥ 0.2 and ΔIC_50_ ≥ 3. Statistical analyses were performed using an unpaired two-sided t-test. Lines represent the median and interquartile range. Data were accessed from the Genomics of Drug Sensitivity in Cancer Drug Combinations dataset(13). **(B)** Combination PARP inhibition and chemotherapy is synergistic in isogenic *Brca-*mutated but not in *Brca-*wildtype cells. ID8*Trp53-/-*, ID8*Trp53-/-Brca1-/-*, ID8*Trp53-/-Brca2-/-* cells were treated with doses of olaparib and carboplatin/paclitaxel for 3 days with vehicle-treated cells as controls. Cell viability was measured using alamarBlue. Synergy analysis was performed using Compusyn, generating a combination index (CI) value where green (<1) indicates synergism, yellow (=1) indicates additivity and red (>1) indicates antagonism. Data are pooled from three biological replicates. **(C)** Colony forming assays of ID8 cells treated with vehicle (DMSO), 1 µM olaparib, 6 µM carboplatin/8 nM paclitaxel (olap+carbo/PTX) for 7 days. Quantification of colony formation is presented as % area normalised to vehicle within each cell line. Statistical analyses were performed using one-way ANOVA. * p < 0.05; **** p < 0.0001; ns = non-significant.

Subsequently, we investigated whether an isogenic model would be suitable to study the effects of PARP inhibitor combination therapies. From assessing synergy, the combination of olaparib and carboplatin/paclitaxel (standard-of-care in ovarian cancer) was antagonistic in *Brca-*wildtype cells (ID8*Trp53-/-*), while the combination therapy was synergistic in ID8*Trp53-/-Brca1-/-* and ID8*Trp53-/-Brca2-/-* cells (Figure 2B). Moreover, the combination therapy was effective in reducing colony growth in *Brca2-/-* cell lines (Figure 2C).

Taken together, this brief report highlighted the importance of exercising caution when using cell line collections in model selection when investigating PARP inhibitor sensitivities and synergies. These data further suggest using isogenic models with specific mutations in *BRCA1* and *BRCA2* and to potentially avoid comparing cell lines from diverse genetic backgrounds, as they may not accurately reflect patient responses.

During this study, Takamatsu *et al*. published a study confirming that cell lines with alterations in *BRCA1* and *BRCA2* had no association with response to PARP inhibitors and platinum agents, as measured by area under the curve (AUC) analysis(20). Using additional measures, such as ‘HRD score’ and mutational ‘Signature 3’, they confirmed that changes associated with homologous recombination deficiency did not demonstrate increased response to PARP inhibitors(20). Additional studies found that signatures related to homologous recombination deficiency had poor predictive power of PARP inhibitor response in colorectal cancer cell lines(21), and *BRCA* status was not predictive of therapy response in a small panel of ovarian cancer cell lines(22). Our collective findings suggest that cell line collections, with diverse genetic backgrounds, do not accurately reflect patient response to PARP inhibitors. Here, we show that this finding extends to synergy studies in that *BRCA1/BRCA2* status in cell line panels is unable to accurately predict the combination effects of PARP inhibitors with chemotherapy.

While we found a positive correlation between *PARP1* expression and response to PARP inhibitors, which is in line with many *in vitro* studies(23, 24), in patients, the relationship between *PARP1* expression and sensitivity to PARP inhibitors is more complicated. Although adequate PARP1 protein is necessary for PARP inhibitor function, it is not clear whether high *PARP1* expression increases sensitivity to PARP inhibitors in patients. Most clinical trials do not investigate *PARP1* expression, although *PARP1* expression correlated with uptake of PARP inhibitors in a small phase I trial(25). Finally, we also revealed that cell lines with lower genome complexity was a better predictor of response to PARP inhibitors than *BRCA* mutations. This suggests that preclinical models with high *PARP1* expression and a simpler genomic profile may be more conducive to identifying clinically translatable effects of PARP inhibitors.

We found that isogenic models with specific mutations in *Brca1* and *Brca2* recapitulated patient sensitivity to PARP inhibitors. These are in line with multiple *in vitro* studies demonstrating the sensitivity of ID8*Trp53-/-Brca1-/-* and ID8*Trp53-/-Brca2-/-* cells to PARP inhibitors, including the original studies that generated the mutant models(10-12). These data translate to the *in vivo* setting, where ID8*Trp53-/-Brca1-/-* and ID8*Trp53-/-Brca2-/-* tumor-bearing mice demonstrated increased survival after rucaparib treatment(12). Furthermore, our data validates that ID8 cells with *Brca* mutations recapitulate patient response to combination PARP inhibitor and chemotherapy(6). While we propose that the ID8 model is useful, particularly due to its ease of use, it is important to note that we utilized a single isogenic cell line. However, these findings have also been observed in other models, such as RPE-1(26). Moreover, while the ID8 model is syngeneic and allows for use in immunocompetent mice, incorporating a representative human model, such as OVCAR-4(27), alongside the ID8 model would help ensure the validity of the results.

Overall, we conclude that standard cell line collections are not the optimal tool to draw conclusions the action of PARP inhibitors or developing new clinical applications. We have shown that PARP inhibitor and chemotherapy combination synergizes to induce reduced viability on an isogenic background, representative of the effects seen in patients. Reverse engineering of other successful combinations applied in patients (eg VEGF inhibitors and PARP inhibitors) could further delineate the most appropriate models for PARP inhibitor research(28).

## STATEMENTS AND DECLARATIONS

### Author contribution

Study conception and design, data collection and analysis, and manuscript preparation were performed by Z.P. and C.E.C. K.J.F. contributed to data collection and analysis. All authors read and approved the final manuscript.

## Acknowledgments

We acknowledge the generosity of funders in supporting our research. Z.P. was supported by the Australian Government Research Training Program Scholarship and a Tour de Cure PhD Support Scholarship (RSP-538-FY2023). Funding for C.E.C. was obtained from the Australian Gynaecological Cancer Foundation (AGCF), assisted by The NSW Government and The Duncan Family, as part of the AGCF Grant 004/01/2024. C.E.C also receives support as recipient of the Lysia O’Keefe Fellowship, and research funding from the Dr Lee MacCormick Edwards Foundation.

## Supplementary Methods

### 1. cBioPortal analysis

Data on *BRCA1, BRCA2* and *PARP* mRNA expression relative to diploid samples, mutation type, copy number alteration status and fraction genome altered were extracted from cBioPortal using the Cancer Cell Line Encyclopedia (CCLE) (Broad, 2019) dataset (Data accessed on 20th February, 2024)(1-3). Data was extracted alongside IC_50_ treatment response to olaparib, talazoparib, rucaparib, veliparib and cisplatin. The oncogenic effect of the variant was annotated using OncoKB™ and Hotspots. “Driver” and variance of unknown significance (“VUS”) mutations were annotated using OncoKB and Hotspots. Cell lines without gene profiling were excluded from the analysis. The list of cell lines along with *BRCA1* and *BRCA2* mutation status can be accessed in Supplementary Tables 1 and 2. To subset ovarian cancer specific cell lines, the search term “_OVARY” was used. Data was plotted and analysed using GraphPad Prism (v10.0).

### 2. Genomics of Drug Sensitivity in Cancer analysis

Data on the synergistic effects of combination olaparib and cisplatin were accessed from the Genomics of Drug Sensitivity in Cancer (GDSC) screening platform(4). Given that ovarian cancer cell lines were not included in this database, we investigated a collection of 46 breast cancer cell lines, as breast cancer is a malignancy where PARP inhibitors have also been approved. Synergy was determined using olaparib as the ‘anchor’ and cisplatin as the ‘library’ where the concentration of olaparib was 10 μM, and cisplatin contained a range doses, the max concentration being 4 μM. According to GDSC, synergy was determined as ≥EMax ≥ 0.2 and/or a ΔIC_50_ ≥ 3. Where there were multiple values for a single cell line, the average was taken. Cell line *BRCA* mutation status and driver annotation was cross-referenced to the CCLE dataset. Cell lines where *BRCA* mutation status could not be determined were excluded from the analysis. All data were downloaded from GDSC Combinations, https://gdsc-combinations.depmap.sanger.ac.uk/.

### 3. Cell culture

*In vitro* experiments were performed using the isogenic ID8 models of ovarian cancer, (RRID: CVCL_IU14). Parental ID8 cells and CRISPR/Cas9-mutant cell lines, ID8*Trp53-/-*, ID8*Trp53-/-Brca1-/-* and ID8*Trp53-/-Brca2-/-* were kindly provided by Prof Iain McNeish (Imperial College London, UK)(5, 6). All cell lines were authenticated and was further confirmed by *Brca1* and *Brca2* Sanger sequencing. Note that a *Trp53* mutant background is utilised because *TP53* mutations are ubiquitous in ovarian cancer(7). Cells were grown in DMEM (Gibco) media supplemented with 4% foetal bovine serum (Sigma-Aldrich HyClone), 1% penicillin/streptomycin (Invitrogen), and 1% insulin-transferrin selenium (Gibco). Cells were routinely passaged when they reached 80% confluency. To passage cells, first, cells were washed with warm phosphate-buffered saline (Gibco), and then incubated with 0.05% Trypsin/EDTA for 5 mins at 37 °C. Culturing media was used to deactivate the trypsin, and cells were transferred to a fresh tissue culture flask. Cells were maintained at 37 °C and 5% CO_2_.

### 4. Drugs

For *in vitro* analyses, cell lines were treated with the following drugs or matched vehicle controls: olaparib (in DMSO; SelleckChem, S1060), carboplatin (in H_2_O; Abcam, ab120828) and paclitaxel (in DMSO; SelleckChem, S1150).

### 5. Cell viability assay

800 cells/well were plated into 96-well plates and media alone was used as a blank. After allowing cells to attach for 8 hours, cells were treated with increasing concentrations (10 pM – 100 μM) of olaparib. After 3 days, alamarBlue reagent (Thermo Fisher Scientific) was added in a 1:10 dilution, and after a 3-hour incubation at 37 °C, fluorescence (excitation 544 nm and emission 590 nm) was measured using the FLUROstar OPTIMA plate reader (BMG LabTech). To calculate cell viability curves, technical replicates were averaged, and the blank (media-only control) was subtracted from sample values. Data was then normalised to solvent control. These values were log transformed, normalised and plotted with a nonlinear fit of least square analysis. IC_50_ values were determined using GraphPad Prism (v10.0).

### 6. Colony forming assay

ID8 cells (300 cells/well) were seeded into 6-well plates as single-cells and left to attach for 6 hours. Cells were then treated with 1 μM olaparib, 6 μM carboplatin/8 nM paclitaxel or the combination of olaparib + carboplatin/paclitaxel and left for 7 days to allow colonies to grow. Media was refreshed once during the 7 days. The colonies were fixed with 16% trichloroacetic acid for 2 hours, washed with distilled water (dH_2_O), and air dried on the bench. Colonies were stained with 0.1% crystal violet for 1 hour before being washed with dH_2_O and left to air dry on the bench. Plates were scanned at 1200 dots per inch using the Perfection V800 Photo scanner (Epson Australia Pty Ltd). The colonies were quantified using ImageJ (v1.0)(8) and the % area was calculated.

### 7. Drug synergy assay

Drug synergy assays were performed based on combination index (CI) method using CompuSyn software (v2.0) (Compusyn, INC). Cells were seeded at 800 cells/well in a 96-well plate and left to attach for 8 hours. Media alone was used as a blank. Based on the IC_50_ of each drug, six drug combinations (three above and three below the IC_50_) were tested to determine the dose-effect curve of olaparib and carboplatin/paclitaxel. Cell viability was measured after 3 days using a 1:10 dilution of alamarBlue and measured using the FLUROstar OPTIMA plate reader. The normalised values were inputted into the CompuSyn software to generate CI values. Microsoft Excel (v16.89.1) was then used to generate a three-colour scale based on the CI values obtained, where synergism (< 1) is represented by green, additive by yellow and antagonism (> 1) by red.

### 8. Statistical analysis

Pearson’s correlation coefficients were calculated using GraphPad Prism (v.10.0). To indicate negative or positive correlations, best-fit linear regressions were fitted onto the graph. All *in vitro* data are presented as the mean ± standard deviation unless otherwise specified. All *in vitro* experiments were performed in biological triplicates. Statistical significance was assessed by unpaired t-tests, two-sided t-test, one-way ANOVA, or two-way ANOVA followed by Tukey’s multiple comparisons test. The specific statistical analysis used for each experiment are provided in the figure legends. P-values < 0.05 were considered as statistically significant.

## Supplementary Figures

**Supplementary Figure 1.**
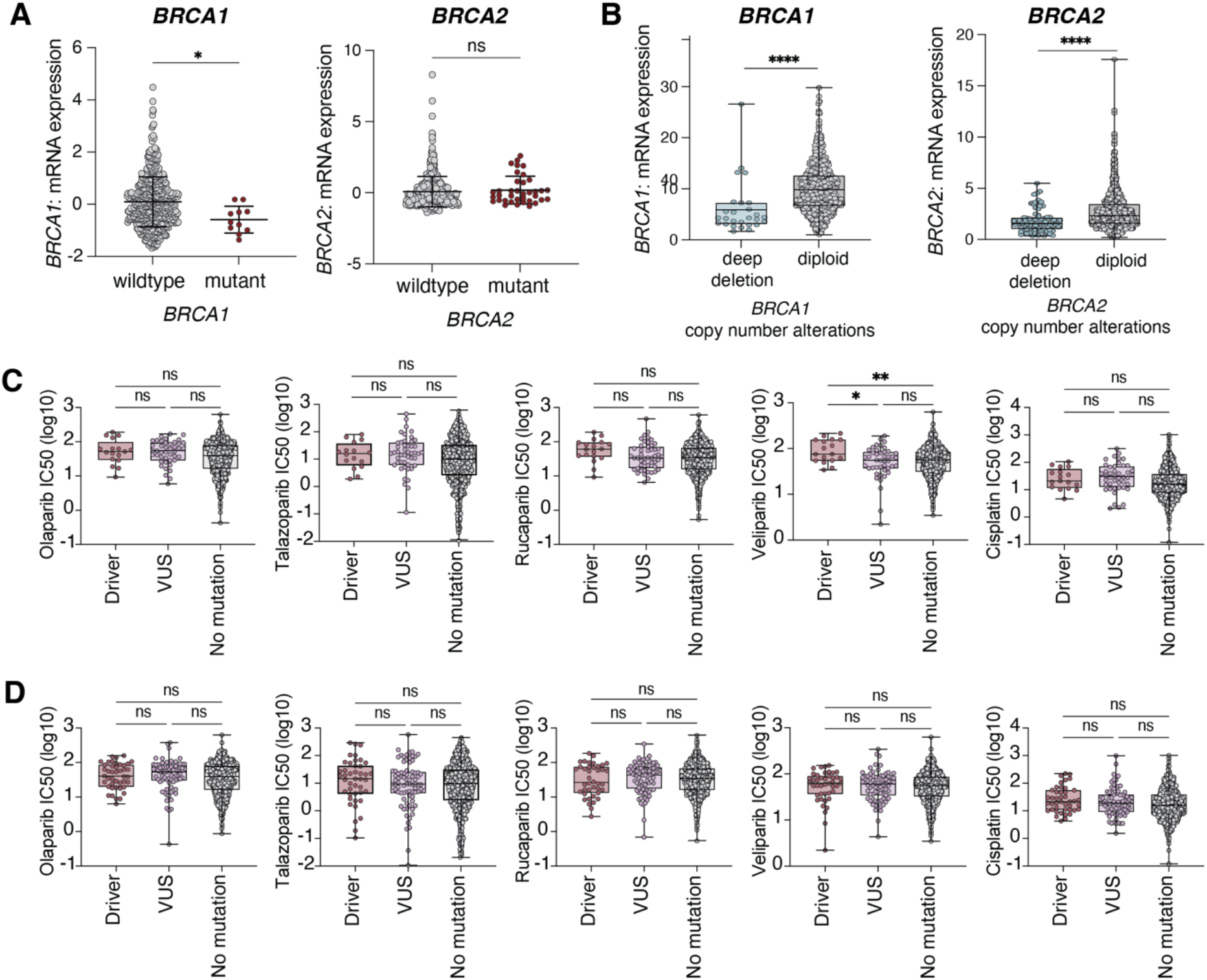
*BRCA1* and *BRCA2* mRNA expression levels and IC_50_ response in cell lines with *BRCA* alterations. **(A)** *BRCA1* and *BRCA2* mRNA expression in wildtype (n = 529 for *BRCA1,* n = 465 for *BRCA2*) and mutant (n = 11 for *BRCA1,* n = 37 for *BRCA2*) cell lines. **(B)** Cell lines with *BRCA1* and *BRCA2* deep deletions exhibit lower mRNA expression levels. *BRCA1* expression is shown for deep deletion (n = 27) and diploid (n = 863) cell lines, while *BRCA2* expression is shown for deep deletion (n = 84) and diploid (n = 818) cell lines. Statistical analyses were performed using an unpaired t-test. **(C)** IC_50_ response of various PARP inhibitors (olaparib, talazoparib, rucaparib, veliparib) and cisplatin in *BRCA1-*mutant cell lines with predicted driver mutations (n = 17-18), variance of unknown significance (VUS) (n = 43-48), and no documented mutations (n = 809-874). **(D)** IC_50_ response to the same treatments in *BRCA2-*mutant cell lines with predicted driver mutations (n = 40-43), VUS (n = 75-81), and no documented mutations (n = 752-819). Statistical analyses were performed using one-way ANOVA. *p < 0.01; **p < 0.001; **** p < 0.0001; ns = non-significant. Data were accessed via cBioPortal.

**Supplementary Figure 2.**
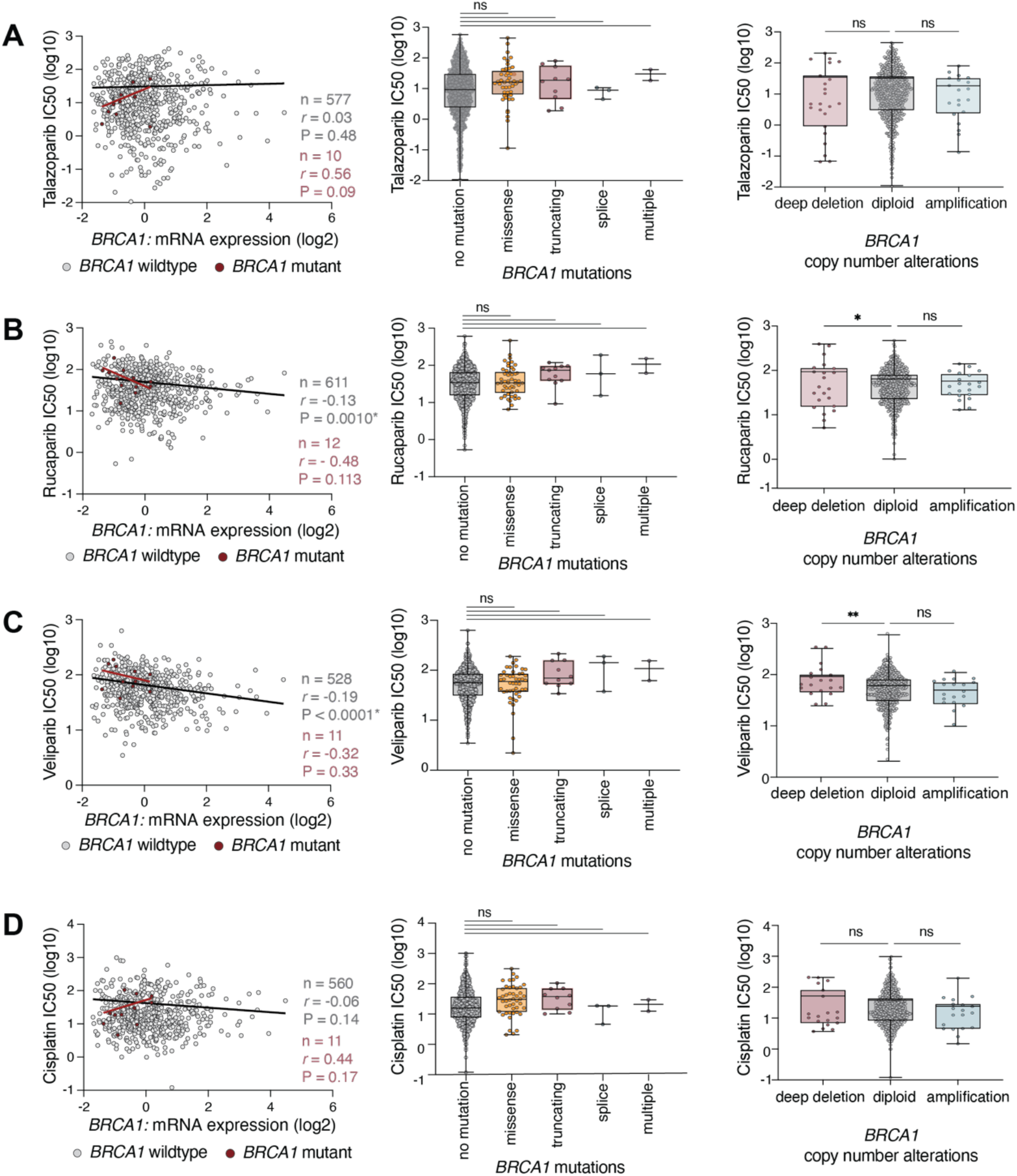
Altered *BRCA1* expression is not associated with an increase in PARP inhibitor and platinum-based chemotherapy sensitivity in cell lines. **(A-D)** IC_50_ responses to PARP inhibitors (olaparib, talazoparib, rucaparib, veliparib) and cisplatin in cell lines with *BRCA1* alterations. Left panels: Correlation between IC_50_ response and *BRCA1* expression relative to diploid samples. Red dots indicate cell lines with predicted driver mutations in *BRCA1* (n = 10-12), while grey dots indicate *BRCA1* wildtype (n = 528-611) cell lines. Best-fit linear regressions (red and black lines) illustrate positive or negative correlations. Middle panels: IC_50_ responses in *BRCA1-*mutant cell lines with different mutation types: no mutation (n = 809-872), missense (n = 46-50), truncating (n = 10-11), splice (n = 3) and multiple (n = 2). Right panels: IC_50_ response in cell lines with different *BRCA1* copy number alteration statuses. IC_50_ values are shown for cell lines classified as deep deletion (n = 19-23), diploid (n = 517-563), and amplification (n = 19-21). Statistical analyses were determined by two-sided Pearson’s correlation test (left panel) and one-way ANOVA (middle and right panels). * p < 0.05; ** p < 0.01; ns = non-significant. Data were accessed via cBioPortal.

**Supplementary Figure 3.**
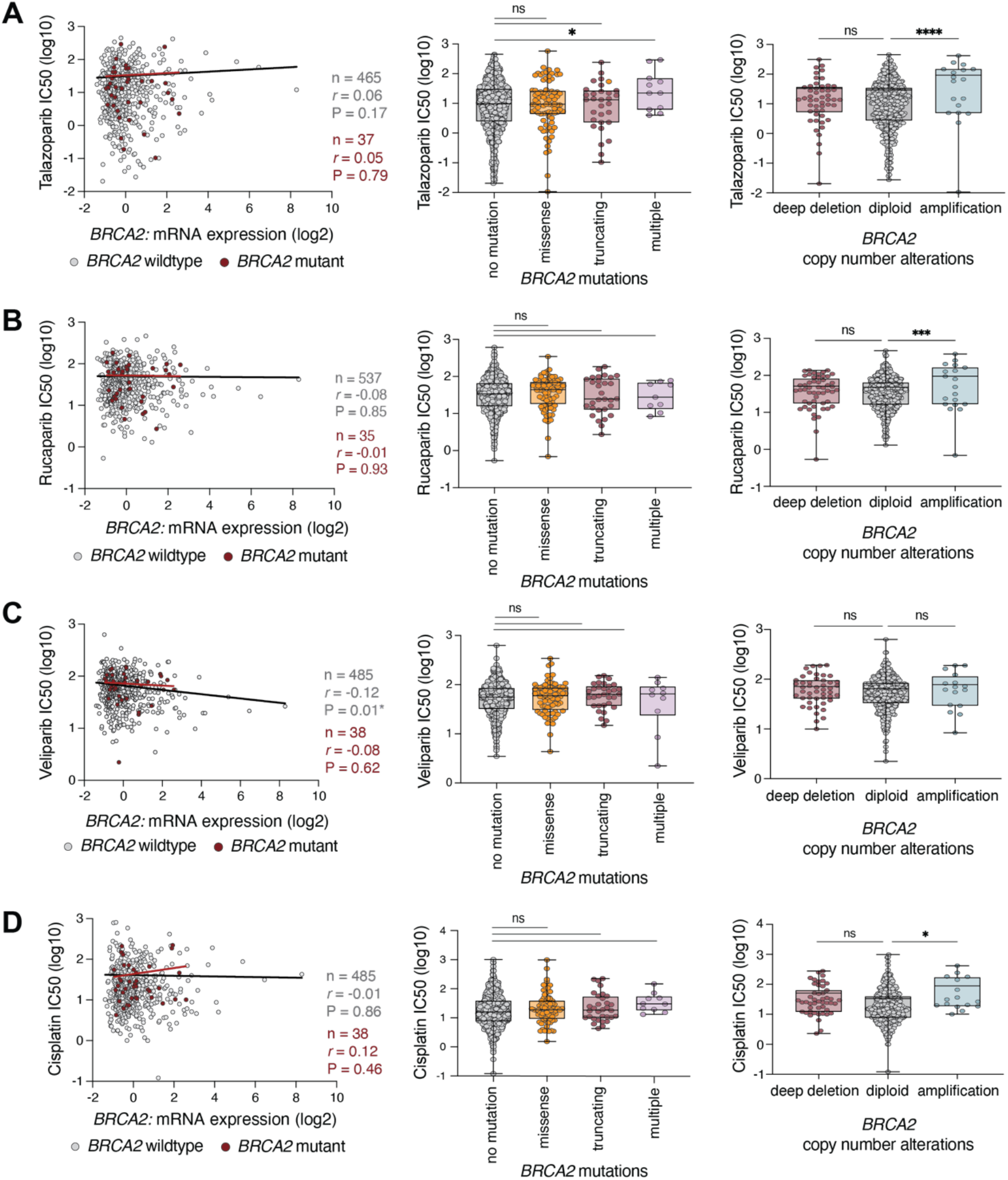
Altered *BRCA2* expression is not associated with an increase in PARP inhibitor and platinum-based chemotherapy sensitivity in cell lines. **(A-D)** IC_50_ responses to PARP inhibitors (olaparib, talazoparib, rucaparib, veliparib) and cisplatin in cell lines *BRCA2* alterations. Left panels: Correlation between IC_50_ response and *BRCA2* expression relative to diploid samples. Red dots indicate cell lines with predicted driver mutations in *BRCA2* (n = 10-12), while grey dots indicate cell lines wildtype *BRCA2* (n = 528-611) cell lines. Best-fit linear regressions (red and black lines) illustrate positive or negative correlations. Middle panels: IC_50_ responses in *BRCA1-*mutant cell lines with different mutation types: no mutation (n = 752-819), missense (n = 76-82), truncating (n = 30-32), and multiple (n = 9-10). Right panels: IC_50_ response in cell lines with different *BRCA2* copy number alteration statuses. IC_50_ values are shown for cell lines classified as deep deletion (n = 47-53), diploid (n = 492-534), and amplification (n = 16-19). Statistical analyses were determined by two-sided Pearson’s correlation test (left panel) and one-way ANOVA (middle and right panels). * p < 0.05; *** p < 0.001; **** p < 0.0001; ns = non-significant. Data were accessed via cBioPortal.

**Supplementary Figure 4.**
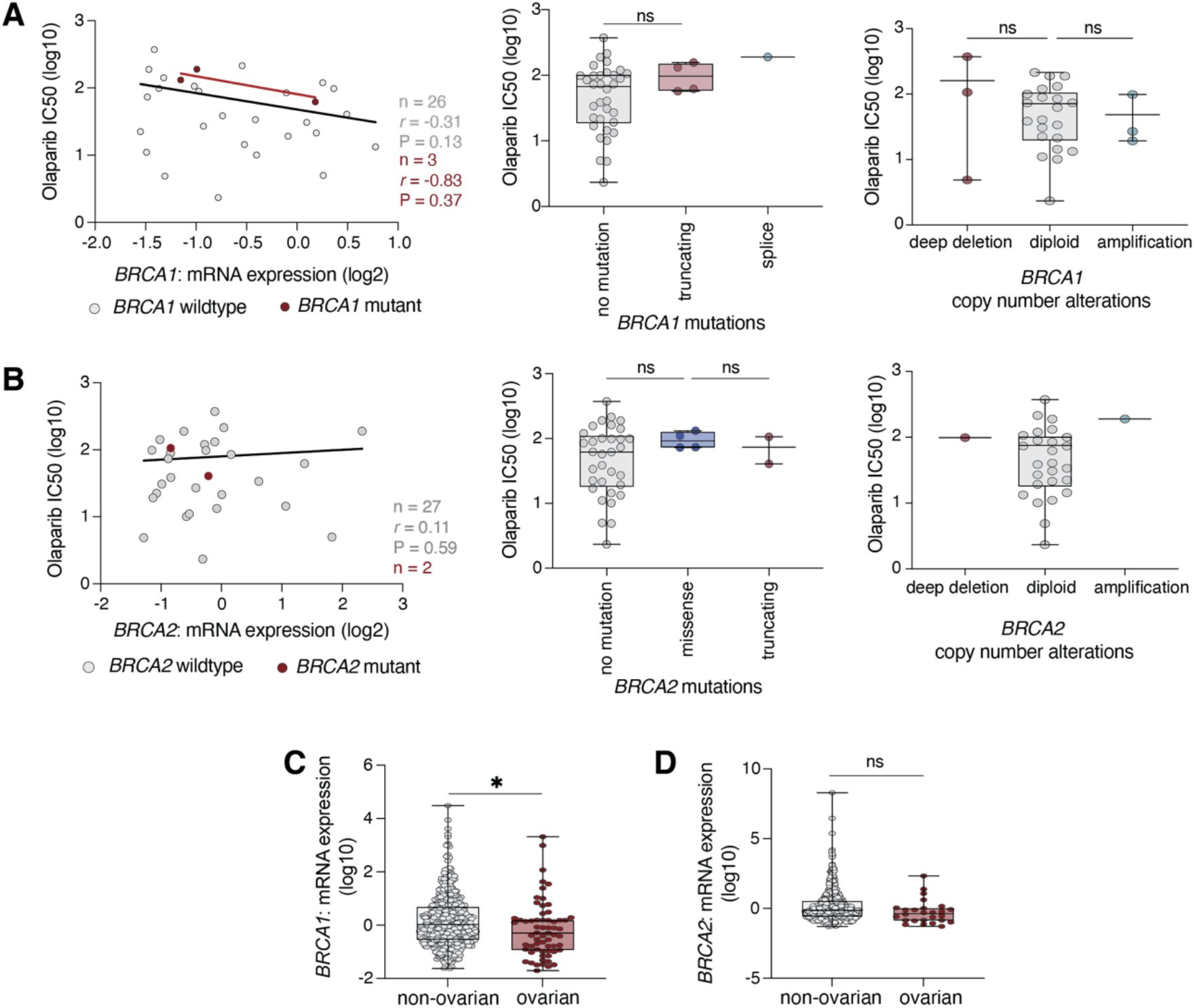
*BRCA1* and *BRCA2* altered ovarian cancer cell line collections are not associated with an increased sensitivity to a PARP inhibitor. **(A-B)** IC_50_ responses to olaparib in ovarian cancer cell lines with **(A)** *BRCA1* and **(B)** *BRCA2* alterations. Left panels: Correlation between IC_50_ olaparib response and *BRCA1/BRCA2* alterations relative to diploid samples. Red dots indicate cell lines with predicted driver mutations in *BRCA1* (n = 3) and *BRCA2* (n = 2), while grey dots indicate wildtype *BRCA1* (n = 26) and *BRCA2* (n = 27) cell lines. Best-fit linear regression (red and black lines) illustrates positive or negative correlations. Linear regression for *BRCA2*-mutant ovarian cancer cell lines were unable to be calculated with small sample size. Middle panels: Olaparib IC_50_ response in *BRCA1-* and *BRCA2*-mutant cell lines with different mutation types: no mutation (n = 34 for *BRCA1,* n = 33 for *BRCA2*), truncating (n = 4 for *BRCA1,* n = 2 for *BRCA2*), splice (n = 1 for *BRCA1*) and missense (n = 4 for *BRCA2*). Right panels: Olaparib IC_50_ response in ovarian cell lines with different *BRCA1/2* copy number alteration statuses. IC_50_ values are shown for cell lines classified as deep deletion (n = 3 for *BRCA1*, n = 1 for *BRCA2*), diploid (n = 22 for *BRCA1,* n = 26 for *BRCA2*) and amplification (n = 3 for *BRCA1,* n = 1 for *BRCA2*). Comparisons for *BRCA2* could not be calculated due to small sample size. Where statistical analyses could be determined, they were by two-sided Pearson’s correlation test (left panels) and one-way ANOVA (middle and right panels). **(C-D)** mRNA expression of **(C)** *BRCA1* and **(D)** *BRCA2* in ovarian (n = 63 for *BRCA1,* n = 26 for *BRCA2*) and non-ovarian cancer (n = 508 for *BRCA1,* n = 508 for *BRCA2*) cell lines. Statistical analyses were determined by unpaired t-test. * p < 0.05; ns = non-significant. Data were accessed via cBioPortal.

